# Data-independent Acquisition-based Proteome and Phosphoproteome Profiling across Six Melanoma Cell Lines Reveals Determinants of Proteotypes

**DOI:** 10.1101/2020.12.14.422682

**Authors:** Erli Gao, Wenxue Li, Chongde Wu, Wenguang Shao, Yi Di, Yansheng Liu

## Abstract

Human cancer cell lines are widely used in pharmacological and systems biological studies. The rapid documentation of the steady-state gene expression landscape of the cells used in a particular experiment may help to improve the reproducibility of scientific research. Here we applied a data-independent acquisition mass spectrometry (DIA-MS) method, coupled with a peptide spectral-library free data analysis workflow, to measure both proteome and phosphoproteome of a melanoma cell line panel with different metastatic properties. For each cell line, the single-shot DIA-MS detected 8,100 proteins and almost 40,000 phosphopeptides in the respective measurement of two hours. Benchmarking the DIA-MS data towards the RNA-seq data and tandem mass tag (TMT)-MS results from the same set of cell lines demonstrated comparable qualitative coverage and quantitative reproducibility. Our data confirmed the high but complex mRNA~protein and protein~phospsite correlations. The results successfully established DIA-MS as a strong and competitive proteotyping approach for cell lines. The data further showed that all subunits of Glycosylphosphatidylinositol (GPI)-anchor transamidase complex were overexpressed in metastatic melanoma cells and identified altered phosphoprotein modules such as BAF complex and mRNA splicing between metastatic and primary cells. This study provides a high-quality resource for calibrating DIA-MS performance, benchmarking DIA bioinformatic algorithms, and exploring the metastatic proteotypes in melanoma cells.

## 1 Introduction

Human cancer cell lines are widely used in biological and biomedical research, serving as an important model system for studying normal and aberrant cellular processes. Comprehensive molecular profiling for multiple cell lines or cell line panels has been demonstrated promising, which connects the genomic alterations to functional networks and pharmacological responses in cancer cells. Just as examples, a pilot study measured the quantitative proteome for 11 common human cell lines and discovered ubiquitous and varying expressions of most proteins ^1^. The Cancer Cell Line Encyclopedia (CCLE) generated multi-layered molecular profiling datasets for 947 human cancer cell lines that encompass 36 tumor types, providing a resource for studying genetic variants, candidate targets, and biological therapeutics in human cancers ^2, 3^. The compilation of CCLE recently was added with a high-quality quantitative proteomics dataset of 375 cell lines using tandem mass tag (TMT) mass spectrometry (MS), which revealed post-transcriptional mechanisms undiscovered by DNA and RNA methods ^4^. At a smaller sample scale, Roumeliotis *et al.* profiled a total of 50 colorectal cancer cell lines with TMT and quantified 9,000 proteins and 11,000 phosphopeptides between cells. This study leveraged a systematic view of proteotype co-variation networks determined by genomic factors in colorectal cancer ^5^. Also, the proteome maps of NCI-60 cell lines were analyzed by different MS techniques ^6-8^.

On the other hand, others and we have reported the instability of cell lines, both genetic and phenotypic, even for the cells of the same name between different laboratories ^9, 10^. For example, in 14 stock HeLa samples from 13 international laboratories, we discovered substantial heterogeneity between the HeLa strains by the total genomic, transcriptomic measurements, as well as the proteomic profiling of 5,000 proteins ^10^. These studies suggested that the previous cell line authentication methods using *e.g.*, short tandem repeat (STR) and single nucleotide polymorphism (SNP) analysis ^11^ might be not sufficient to document the particular cell line used in an experiment, prompting rapid documentation of precise gene expression status of the cells ^10^. Although transcriptomics can effectively record the basic state of the cells, proteins are the executors of the function encoded by a cell’s genome. The direct proteomic measurement should be alternatively considered for documenting the cell’s molecular landscape.

As above, there are pressing needs for both high-throughput characterization of multiple cells and proteomic documentation of individual cells. Furthermore, because dynamic phosphorylation plays a major role in regulating many cellular processes, phosphoproteomic profiling was recently used in defining the proteome “activity” in cancer cell line panels ^8^. Together, it is imperative to establish a fast, cost-effective, reproducible method for profiling the cell “proteotype” ^12^, ideally through both proteome and phosphoproteome profiling. Previously, the integrative proteomic and phosphoproteomic measurements have been successfully performed by coupling peptide-fractionation to either shotgun-or TMT-MS workflows in cancer cell lines and clinical tissues ^5, 8, 13-16^. The data-independent acquisition mass spectrometry (DIA-MS) ^17-19^ provides an alternative, simple, and robust MS workflow for profiling proteotype through the concurrent proteomic and phosphoproteomic analysis ^20^, without the need of extensive peptide-level fractionation and the costly tagging reagents. However, DIA-MS was previously acknowledged to generate 15-20% less protein identifications than TMT workflow with fixed instrument time ^21^. Besides, very few studies have analyzed both proteome and phosphoproteome of cell lines by DIA-MS. Nevertheless, the complex quantitative relationship among transcripts, proteins, and phosphosites has not been characterized in a standard DIA-MS dataset previously. Herein, we describe a high-quality DIA dataset acquiring single-shot measurements for both proteome and phosphoproteome of six metastatic and primary melanoma cell lines. We then assessed the qualitative and quantitative features of this DIA dataset and systematically investigated the mRNA-protein and proteome-phosphoproteome correlations. We further discussed the appropriate phosphoproteomic normalization strategies using the plentiful peptide-level identifications in DIA. This work provides a valuable resource for evaluating DIA-MS performance and understanding the melanoma cell proteotypes.

## 2 Materials and methods

### 2.1 Cell culture

The metastatic melanoma cancer cells (ATCC TCP-1014) and the primary melanoma cancer cells (ATCC TCP-1013) were purchased from ATCC. The three metastatic cell lines include RPMI-7951 (ATCC HTB-66, named “**7951**” hereafter), SH-4 (ATCC CRL-7724, named “**SH4**”), and SK-MEL-3 (ATCC HTB-69, named “**HTB69**”). The three primary cell lines include SK-MEL-1 (ATCC HTB-67, named “**SK**”), A375 (ATCC CRL-1619, named “**A375**”), and G-361 (ATCC CRL-1424, named “**G361**”). The routine cell culture protocol was detailed previously^10^. In brief, cells were cultured in 5% CO_2_ and 37° in either DMEM (#10564011, for 7951, HTB69, A375 and G361 cells) or RPMI Medium (#72400047, for SH4 and SK cells) supplemented with 10% FBS (Sigma Aldrich), together with a penicillin/streptomycin solution (Gibco). Cells were harvested at 80% confluence for mRNA and protein extractions.

### 2.2 RNA extraction, quality control, library preparation, and sequencing

Cells were washed by PBS twice, snap-frozen, and then lysed with the QIAShredder columns (Qiagen) according to manufacturer’s instructions. Total RNA was isolated with the Qiagen RNeasy Mini Kit (#74104, QIAGEN), including DNA digestion step using the RNAase-Free DNAase set kit (#79254). RNA samples were quantified and checked for quality control using the Agilent 4200 Tapestation RNA Screentape assay. Samples with RINs greater than 7 were selected for library preparation. Library preparation was performed using the KAPA Biosystems mRNA HyperPrep Kit, in which samples were normalized with a Total RNA input of 1000 ng and library amplification with 8 PCR cycles. Libraries were validated using the Agilent 4200 Tapestation D1000 assay and quantified KAPA Library Quantification Kit for Illumina® Platforms kit. Sequencing was done on an Illumina NovaSeq 6000 using the S4 XP workflow. Libraries were pooled to 1.25% in order to achieve 25M read pairs for each library. Three dish replicates per cell line were used for RNA sequencing. A simple TPM (Transcripts Per Kilobase Million) cutoff ^22^ of 0.1 was applied to retain possibly expressed genes at the transcriptomic level.

### 2.3 RNA data procession and analysis

The reads were trimmed to remove low-quality based-calls, and the Minimum accepted length was 45 bases. If the trimming reduces the read length below 45 bases, that read is discarded. We used HISAT2 ^23^ for alignment of the trimmed reads to the reference genome hg38, with GENCODE annotation ^24^. We then used the StringTie /Ballgown ^25^ to generate gene counts and transcript abundance estimates from the alignments. TPMs reported were log2-transformed for statistical analysis.

### 2.4 Protein extraction and digestion

Cultured cells were harvested and digested as previously described ^26, 27^. Briefly, cells were washed three times by PBS, harvested, and snap-frozen. The cell pellets were then lysed by adding 10 M urea containing complete protease inhibitor cocktail (Roche) and Halt™ Phosphatase Inhibitor (Thermo), and were further ultrasonically lysed at 4 °C for 2 min using a VialTweeter device (Hielscher-Ultrasound Technology), and centrifuged at 18,000 × g for 1 hour to remove the insoluble material. A total of 800 μg supernatant proteins (determined by BioRad Bradford assay) were reduced by 10 mM tris-(2-carboxyethyl)-phosphine (TCEP) for 1 hour at 37 °C and 20 mM iodoacetamide (IAA) in the dark for 45 min at room temperature. Five volumes of precooled precipitation solution containing 50% acetone, 50% ethanol, and 0.1% acetic acid were added to the protein mixture and kept at −20 °C overnight. The mixture was centrifuged at 18,000×g for 40 min. The precipitated proteins were washed with 100 % acetone and 70% ethanol with centrifugation at 18,000×g, 4°C for 40 min. Following that, 300 μL of 100 mM NH_4_HCO_3_ was added in to each sample with sequencing grade porcine trypsin (Promega) at a ratio of 1:20 overnight at 37 °C. The resulted peptide mixture was acidified with formic acid and then desalted with a C18 column (MarocoSpin Columns, NEST Group INC) following manufacturer’s instructions. The amount of the final peptides was determined by Nanodrop (Thermo Scientific). Duplicate dishes per cell line were used for proteomics analysis.

### 2.5 Phosphoproteomics sample preparation

The phosphopeptide enrichment was performed using the High-Select™ Fe-NTA kit (Thermo Scientific, A32992) according to the manufacturer’s instructions ^15^. Briefly, the resins of spincolumn were aliquoted, incubated with 200 μg of total peptides (see above) for 30 min at room temperature, and transferred into the filter tip (TF-20-L-R-S, Axygen). The supernatant was then removed by centrifugation. Then, the resins adsorbed with phosphopeptides were washed sequentially with 200 μL × 3 washing buffer (80% ACN, 0.1% TFA) and 200 μL × 2 H_2_O to remove nonspecifically adsorbed peptides. The phosphopeptides were then eluted off the resins by 100 μL × 2 elution buffer (50% ACN, 5% NH_3_•H_2_O). The centrifugation steps were all kept at 500 g, 30 sec. The eluates were dried by speed-vac and stored in −80 °C before MS measurements.

### 2.6 DIA-MS measurement

The peptide samples were resolved in 2% ACN, 0.1% FA, and 1 μg of peptides or enriched phosphopeptides was injected per each single MS injection. The DIA-MS measurement was performed mainly as described ^28^. Briefly, LC separation was performed on EASY-nLC 1200 systems (Thermo Scientific, San Jose, CA) using a 75 μm × 50 cm C18 column packed with 100A C18 material. A 120-min measurement with buffer A (0.1% formic acid in H_2_O) and buffer B (80% acetonitrile containing 0.1% formic acid) mixed and configured as below to elute peptides from the LC: Buffer B was increasing from 6% to 37% in 109 mins, increased to 100% in 3 mins, and then kept at 100% for 8mins. The flow rate was kept at 300 nL/min with the temperature-controlled at 60 °C using a column oven kit (PRSO-V1, Sonation GmbH, Biberach, Germany). The Orbitrap Fusion Lumos Tribrid mass spectrometer (Thermo Scientific) instrument coupled to a nanoelectrospray ion source (NanoFlex, Thermo Scientific) was used as the DIA-MS platform for both proteomic and phosphoproteomic analyses. Spray voltage was set to 2,000 V and heating capillary temperature at 275 °C. All the DIA-MS methods consisted of one MS1 scan and 40 MS2 scans of variable isolated windows ^28^, with 1 m/z overlapping between windows. The MS1 scan range is 350 - 1650 m/z, and the MS1 resolution is 120,000 at m/z 200. The MS1 full scan AGC target value was set to be 2.0E5, and the maximum injection time was 100 ms. The MS2 resolution was set to 30,000 at m/z 200 with the MS2 scan range 200 - 1800 m/z, and the normalized HCD collision energy was 28%. The MS2 AGC was set to be 5.0E5, and the maximum injection time was 50 ms. The default peptide charge state was set to 2. Both MS1 and MS2 spectra were recorded in profile mode.

### 2.7 Proteomics and phosphoproteomics data procession and analysis

DIA-MS data analyses for proteomics and phosphoproteomics were performed using Spectronaut v14 ^29, 30^, both with the “DirectDIA” pipeline (*i.e.*, an optimal spectral library-free pipeline ^31^). This means the DIA runs were all directly searched against Swiss-Prot protein database (September 2020, 20,375 entries). For the identification of the total proteomic dataset, the possibilities of Oxidation at methionine and Acetylation at the protein N-terminals were set as variable modifications, whereas Carbamidomethylation at cysteine was set as a fixed modification. For the “DirectDIA” database searching on the phosphoproteomic dataset, in addition to the above peptide modification settings, the possibility of Phosphorylation at serine/threonine/tyrosine (S/T/Y) was enabled as a variable modification. Overall, both peptide- and protein-FDR (based on Qvalue) were controlled at 1%, and the data matrix was filtered by Qvalue. In particular, the PTM localization option in Spectronaut v14 was enabled to locate phosphorylation sites ^32,33^ for the entire phosphoproteomic experiment, with the PTM score >0.75 ^33^ applied in at least one of the twelve single-shot phosphoproteomic injections, generating Class-I sites ^34^ for all phosphopeptides at the whole experiment level. The PTM score of 0 was used for estimating and reporting the total number of identified phosphosites (that may not be all localized) and for accepting quantitative values in each sample for all Class-I phosphosites identified at the experiment level. All the other Spectronaut settings for identification and quantification were kept as default, meaning that *e.g.*, the “Inference Correction” was enabled, the “Global Normalization” (on “Median”) was used, the quantification was performed at the MS2 level using peak areas, and the Top 3 peptide precursors (“Min: 1 and Max: 3”) were summed for representing protein quantities in all DIA analyses. For each localized phosphosite, the corresponding phosphopeptide precursors with the least missing values were taken for quantification between samples. The quantitative peak areas for protein and phosphopeptides were then log2-transformed for downstream statistical analysis.

### 2.8 Bioinformatics

The biological gene list annotation and enrichment analysis incorporating Cytoscape^35^ and MCODE based analysis ^36^ were performed by Metascape ^37^ (https://metascape.org/) using the “Multiple Gene List” function with default statistical cutoffs. To embrace as many genes as the input for studying the biology of melanoma metastasis, the simple statistical student’s *t-test* was used and P<0.05 was set as a cutoff. Pearson and Spearman correlation coefficients were calculated using R (functions cor() or cor.test() to infer statistical significance). The colored scatterplots from blue-to-yellow were visualized by the “heatscatter” function in R package “LSD” using a twodimensional Kernel Density Estimation. Online consensus Survival webserver for Skin Cutaneous Melanoma (OSskcm, http://bioinfo.henu.edu.cn/Melanoma/MelanomaList.jsp)^38^ was used to estimate the survival outcome of GPAA1 mRNA expression using the data source of “Combined” or “TCGA” options with the patients split by “Upper 30% VS Lower 30%”. The heatmap following hierarchical clustering analysis (HCA) was created using the R package “pheatmap”. The principal component analysis (PCA) was performed using R function prcomp(). GraphPad Prism (v9) was used to generate the histogram and scatter plots for individual columns.

### 2.9 Data availability

RNA-seq data were uploaded to GEO repository and are available on GEO (GSE162270, data will be available upon paper publication).

The mass spectrometry proteomics data have been deposited to the ProteomeXchange Consortium via the PRIDE partner repository ^39^ with the dataset identifier PXD022992 (data will be available upon paper publication).

## 3 Results and discussion

### 3.1 Single-shot DIA-MS achieving exquisite sensitivity in profiling proteome and phophosproteome

To establish a cost-effective proteotyping method that is generally applicable for individual cancer cell lines, we applied a workflow incorporating our single-shot 2-hour DIA-MS method ^28^ and an improved, spectral library-free ^31^ DirectDIA algorithm (see **Methods**). We quantitatively profiled the cell proteotype via both proteome and phosphoproteome on six melanoma cell lines, including 7951, HTB69, and SH4 cells that are metastatic and A375, SK, and G361 cells that are nonmetastatic. Our DIA-MS workflow was able to detect 8,110 ± 31 protein groups corresponding to 90,588 ± 687 unique peptides in each single-shot of these cell lines, with both peptide- and protein-FDR strictly controlled below 1% (**Figure 1 and Supplementary Table S1**). This analytical coverage represents almost 80% of the total proteome expressed in a cancer cell line under a given condition ^40^. Furthermore, on average, 39,808 unique phosphopeptides were identified in each MS shot under the same FDR threshold. Impressively, high sensitivity was consistently achieved between different cell lines and MS injections - the detection rate variation was merely ~0.5% for the total proteome profiling and ~3% for the phosphoproteome measurement (**Figure 1**). These numbers therefore suggest that DIA-MS stably identified substantial proteins and phosphopeptides among the six cell lines without the need of building spectral libraries.

**Figure 1.**
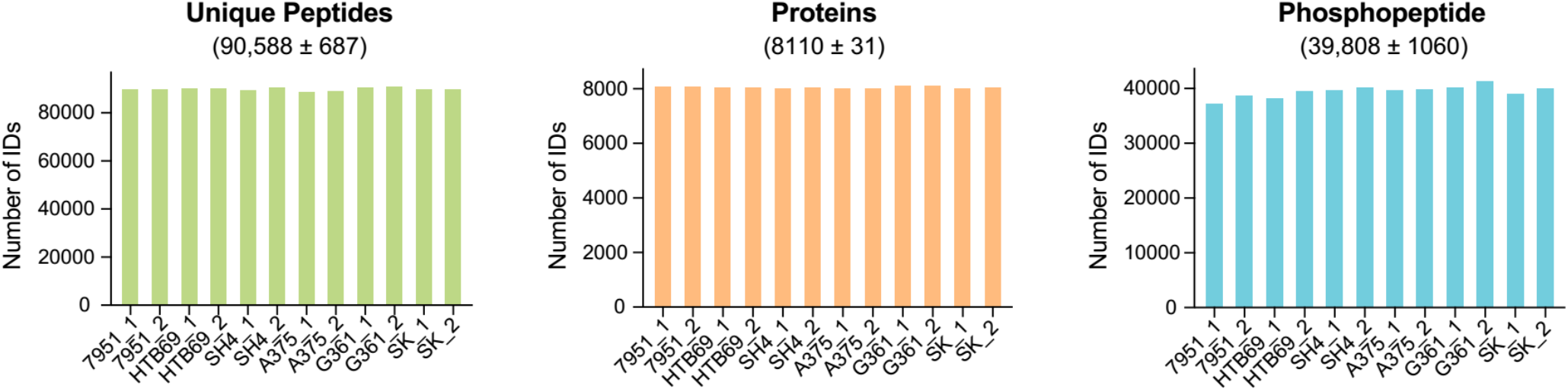
The numbers of unique peptides (left panel), protein groups (middle panel), and phosphopeptides (right panel) identified in the single-shot MS measurement of the six melanoma cell lines. A 2-hour measurement time was adopted for each DIA-MS. Both peptide and protein FDR were controlled at 1%.

### 3.2 Benchmarking the proteomic and phosphoproteomic profiling by RNA-seq analysis

We further assessed the performance of DIA-MS based proteotyping by comparison to transcriptomic profiling, which is relatively more developed than proteomic profiling. Based on RNA sequencing (RNA-Seq), a total number of 13,527 genes could be profiled, in comparison to 8,435 and 6,417 genes respectively covered by proteomics and phosphoproteomics in the same cells (**Figure 2A**). A total number of 7,560 genes was profiled with both RNA and protein expression values, which compares favorably to previous studies ^41, 42^. Compared to total proteomics, the phosphoproteomics covered an extra list of 1,416 genes, but did not detect any phosphosite for 3,434 genes. This result suggests that the phosphorylations of around 40% of the proteome (*i.e.*, 3,434/ 8,435 proteins) might be of extremely low stoichiometry or even not existing. Collectively, nearly 10,000 genes were measured by proteomics and phosphoproteomics (**Figure 2A**), with a total MS measurement time of only 4 hours.

**Figure 2.**
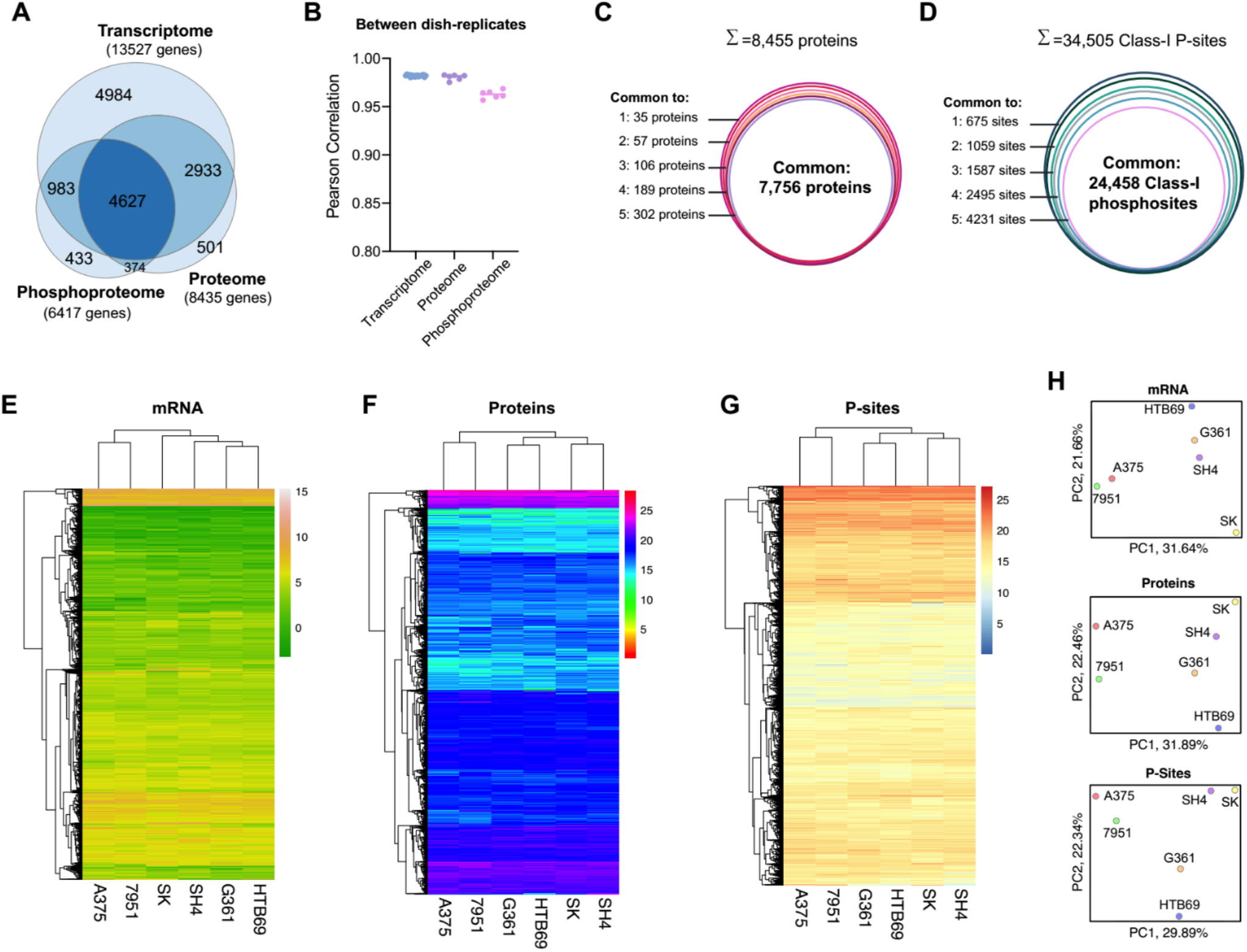
Benchmarking the proteomic and phosphoproteomic results with the RNA-seq data. (A) The Venn diagram between the measured proteome, phosphoproteome and transcriptome of the cell lines. (B) Pearson correlation between dish-replicates, grouped by transcriptome, proteome and phosphoproteome. (C) The number of overlapped proteins measured across the six cell lines. (D) The number of overlapped Class-I phosphosites measured across the six cell lines. (E) Hierarchical clustering analysis of the transcript profiles in the six cell lines. (F) Hierarchical clustering analysis of the protein profiles measured in the six cell lines. (G) Hierarchical clustering analysis of the phosphopeptide (P-site) profiles measured in the six cell lines. (H) Principal component analysis of mRNA, protein and P-site profiles.

Besides the detection performance that is close to RNA-Seq, the reproducibility of quantification represents another significant highlight of DIA-MS. We correlated the TPM values or the DIA-MS peak areas for mRNA, protein, and phosphopeptides between dish-replicates for each cell line; and we found that the dish-replicates were always clustered together (**Supplementary Figure S1**). DIA-MS achieved a replicate-correlation as excellent as RNA-Seq (*i.e.*, Pearson R=0.98 for RNA data, while R= 0.98 and 0.96 for protein and phosphopeptide abundances, **Figure 2B**). Following the multicell line comparison performed by Geiger *et al.* ^1^, we consolidated the quantitative data for 7,756 proteins and 24,458 Class-I phosphosites (P-sites) ^34^ across all the six melanoma cell lines without any missing values (**Figure 2C-D**). Because of the exemplary quantitative reproducibility, the greater reduction of phosphosites across cell lines than the protein identities (*i.e.*, from 34,505 to 24,458 versus from 8,455 to 7,756, **Figure 2D**) could be ascribed to the larger biological variation between phosphoproteomes than between proteomes (**Supplementary Figure S1C and S2**). The multi-omics datasets could be used to discern the global similarity between the six cell lines by hierarchical clustering analysis (HCA, **Figure 2E-F**) and principal component analysis (PCA, **Figure 2H**). In both HCA and PCA, while the 7951 and A375 cells formed a group throughout the mRNA, proteomic, and phosphoproteomic profiling, the other four cell lines (*i.e.*, SK, SH4, G361, and HTB69) were consistently clustered. Notably, this global pattern did not recapitulate the metastatic and nonmetastatic property of melanoma cells, suggesting that the drastic individual genomic variability, rather than the metastatic tumor signatures, dedicated the global clustering results.

In summary, our results suggest that DIA-MS based proteotyping reached a close coverage and an equally great quantitative reproducibility compared to RNA-Seq measurement.

### 3.3 Comparing DIA-MS results to a state-of-the-art TMT dataset for cellular proteomic profiling

The newly added proteomic dataset to the Cancer Cell Line Encyclopedia (CCLE) presents a landmark resource to the community, quantifying an average of 9,175 proteins for 375 cancer cells ^4^. To generate this CCLE dataset, the authors utilized the combination of 10-plex TMT labeling (TMT10), deep peptide-level fractionation, and the synchronous precursor selection (SPS)-based MultiNotch MS3 technique which improves the quantification accuracy in TMT-workflows ^43^. In a recently updated workflow described by the same group, the high-Field Asymmetric Ion Mobility Spectrometry (FAIMS), realtime database searching, and the 16-plex TMTpro labeling were integrated, and a similar proteome coverage was achieved for eight cell lines ^44^. Because the CCLE-TMT dataset ^4^ contains four of six melanoma cell lines we measured in the present study (i.e., 7951, HTB69, SH4, and A375), we benchmarked our DIAMS data to TMT results accordingly. We firstly compared the identification performance based on the unique gene identities. We found that DIA-MS measured the proteome for 7,790 genes *without any missing values* among the four cell lines, whereas the CCLE-TMT measured 7,398 genes, 5.0% less than DIA-MS (**Figure 3A**). However, a further inspection of CCLE-TMT data indicated that 7951 and SH4 cells were measured in the same TMT10 batch, whereas A375 and HTB69 cells were measured in the other two 10-plex batches. Interestingly, when it comes to 7951 and SH4 only, the CCLE-TMT reported the relative quantification for 8,719 genes, denoting a 9% increase compared to DIA-MS (**Figure 3B**). The nearly 10% increase seems to be fairer for TMT, because the TMT would normally accommodate a small number of cell lines in only one experimental batch. Taken together, the above comparisons indicate that ***a)*** the high-quality DIA-MS still identifies about 9-10% fewer proteins than those in the TMT approach in one batch, and ***b)*** even with the proper “bridging sample”, the large-scale TMT profiling using multiple batches could modestly or moderately impact the final quantification outcome between samples. Finally, we checked the relative fold-change between 7951 and SH4 quantified by TMT or DIA-MS, respectively. Using the data from the overlapping 6,900 proteins, we determined the cross-approach correlation to be strong (R=0.67, Pearson correlation, **Figure 3C**), despite that the cells were cultured in different labs and independent proteomic protocols were used.

**Figure 3.**
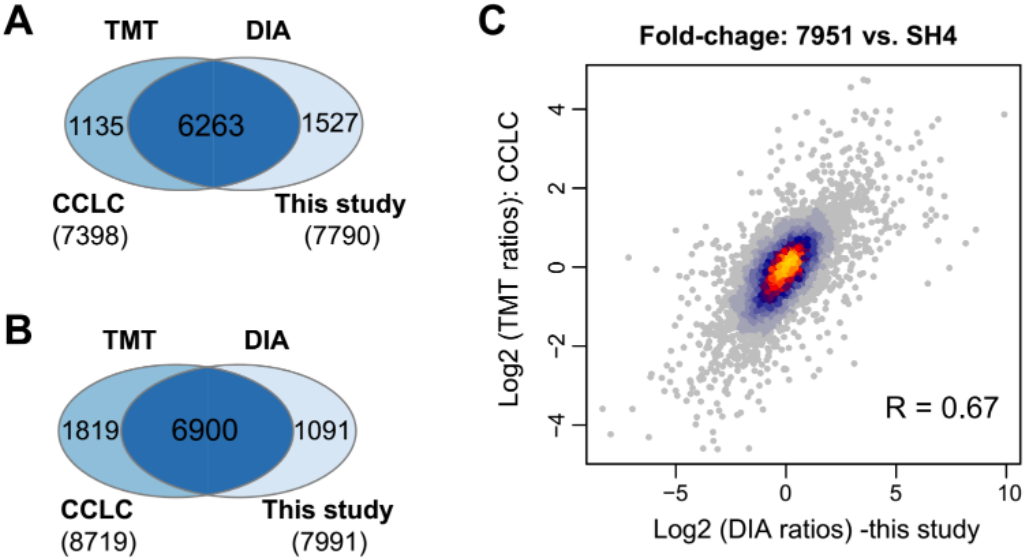
Benchmarking DIA-MS results in this study with the TMT data in Cancer Cell Line Encyclopedia (CCLE). (A) The overlapped proteins without any missing values measured by DIA-MS and TMT. (B) The overlapped proteins of SH4 and 7951 cells measured by DIA-MS and TMT. Note these two cell lines were measured in the same TMT-plex batch. (C) The scatter plot of the protein fold changes between SH4 and 7951 cells measured by DIA-MS and TMT.

To conclude, our data suggest that high-quality DIA-MS could achieve qualitative and quantitative results that are both comparable to TMT. It should be stressed, however, that our small-scale comparison here does not aim to provide a systematic comparison between DIA-MS and TMT - Both approaches have respective advantages over the other. For example, DIA-MS could be more flexible when only one or two cell lines used in an experiment need to be proteotyped ^10^ or when the absolute label-free quantification is desired ^45^, whereas TMT workflow could save loading amounts for precious samples due to its pooling strategy. For a more completed comparison between DIA-MS and TMT with the same instrument time, please refer to Muntel *et al.* ^21^

### 3.4 Quantitative relationships of mRNA vs. protein and proteome vs. phosphoproteome

The quantitative relationship between mRNA and protein has been extensively discussed ^41, 42^. While the absolute mRNA~protein correlation analysis (normally performed within one sample) could be affected by the across-gene variation, the relative mRNA~protein correlation (performed between samples and conditions) can remove this confounding factor for inferring the significance of the post-transcriptional regulation ^41, 42, 46, 47^. The perspective of absolute and relative relationships may also provide insights for analyzing regulations at other molecular layers ^48^. Herein, our well-matched mRNA~protein and protein~phosphosite datasets allow for a rigorous assessment of the variability of protein expression and phosphorylation in the genomic context of steady-state melanoma cell lines. *First*, we correlated the mRNA TPM values and DIA-MS readouts in the log-scale for all the ~7500 genes in each cell line (**Figure 4A, upper panel**). The absolute correlation was 0.58-0.64 (Spearman correlation coefficient ρ) with an average of 0.61, consistent with previous high-quality datasets ^42^. *Second*, we compared the fold-change of mRNA and protein values of each cell line to the *averaged* values across six cells so that a relative correlation could be inferred in each cell line (**Figure 4A, lower panel**). This analysis yielded correlations of 0.54-0.66, with an average of 0.61 as well. The high mRNA~protein correlations at both relative and absolute scales (**Figure 4B**) agree with the previous notion that mRNA levels primarily contributes to protein concentrations in the steadystate ^41, 42^. *Third*, to uncover Protein~P-site relationship, we performed similar correlation analyses (**Figure 4C**). The absolute Protein~P-site abundance correlations were quite low (ρ= 0.12-0.18), compared to the relative correlations (ρ= 0.49-0.53, **Figure 4D**). The absolute correlation here could be affected by the abundance differences among individual P-sites of the same protein, the flyability in MS difference between phosphopeptide and non-phosphopeptide, and intrinsic, variable P-site/Protein ratios. Therefore, the relative analysis reflected that many of phosphoproteome abundance regulations conceivably followed the proteome level.

**Figure 4.**
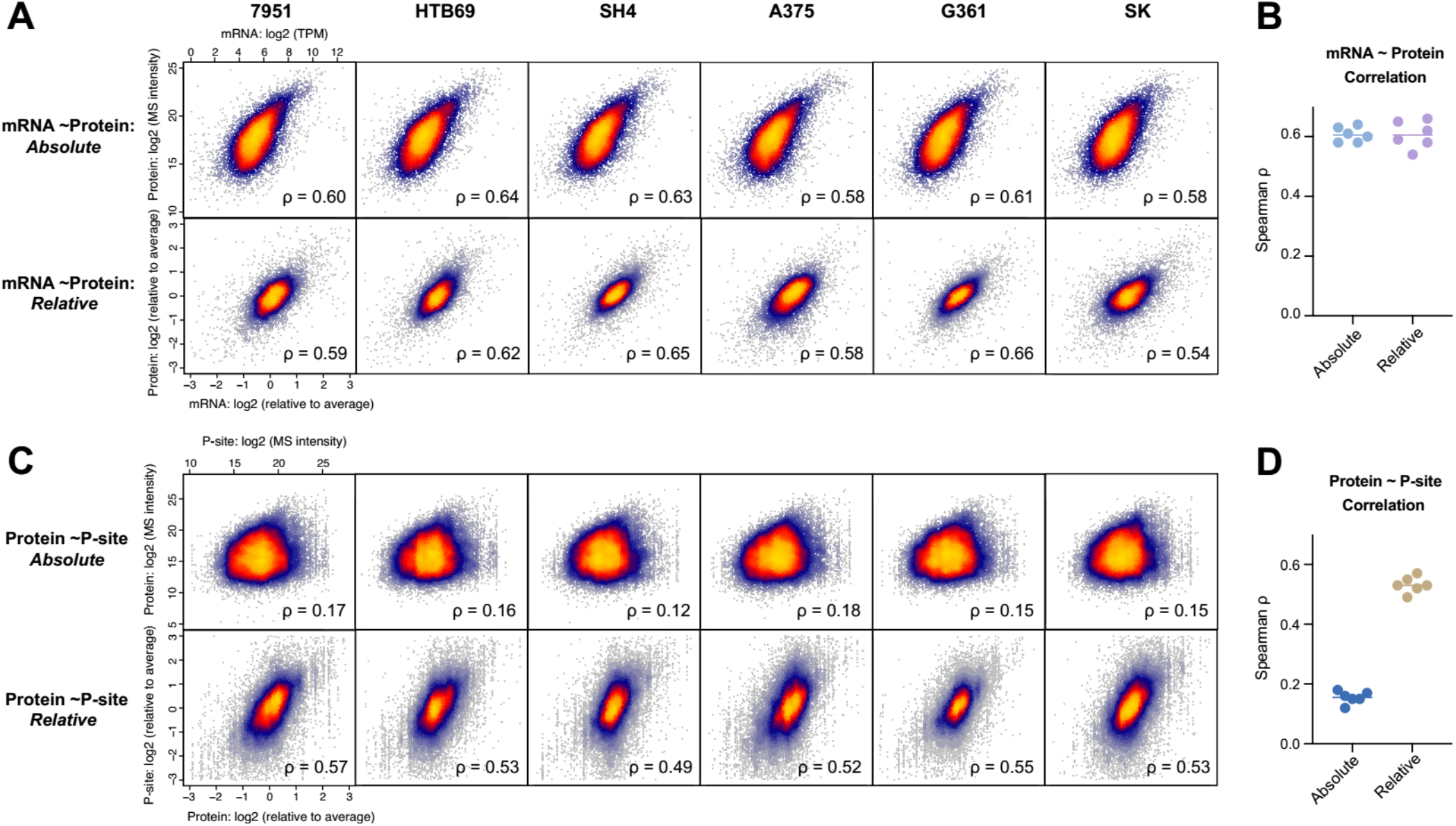
Absolute and relative quantification relationships of mRNA vs. Protein and Protein vs. phosphosite. (A) The scatter plot of mRNA and protein quantities in the absolute (upper panel) and relative (lower panel) scales, separated by six individual six cells. (B) The Spearman correlation between mRNA and protein summarized. (C) The scatter plot of protein and phosphosite quantities in the absolute (upper panel) and relative (lower panel) scales, separated by six individual six cells. (D) The Spearman correlation between protein and phosphosite summarized.

Besides the overall mRNA-protein correlation, the genespecific correlation analysis was demonstrated to be powerful in associating the different genes and pathways with the extent of post-transcriptional regulation among samples ^14, 46, 49, 50^. Thus, we distributed the gene-specific mRNA~protein and Protein~P-site correlation coefficients. We then functionally annotated those genes with low correlations (ρ <0.2, **Figure 5**). From the low mRNA~protein correlations, we discovered that those genes enriched in *e.g.*, Cell cycle, DNA repair, Mitochondrion organization, Autophagy, Respiratory electron transport, and other regulatory processes (-Log10_P value >10, based on Metascape ^37^, **Figure 5A**) were regulated significantly at the post-transcriptional level. On the other hand, the phosphorylation events enriched in processes such as mRNA processing, Cell cycle, Nuclear transport, Covalent chromatin modification, Cellular response to stress, and Membrane trafficking tend to be regulated independently on the corresponding protein abundance (-Log10_P value >20, **Figure 5B**), because they preferred to harbor low Protein~P-site correlations. Thus, although the sample size is small (n=6 cell lines), our preliminary analysis indicates protein-level remodeling and phosphorylation modification can be remarkably different for various pathways and biological processes. The results also urge the multi-layered, with-in gene variance analysis to be performed in more cell lines and cell panels in the future.

**Figure 5.**
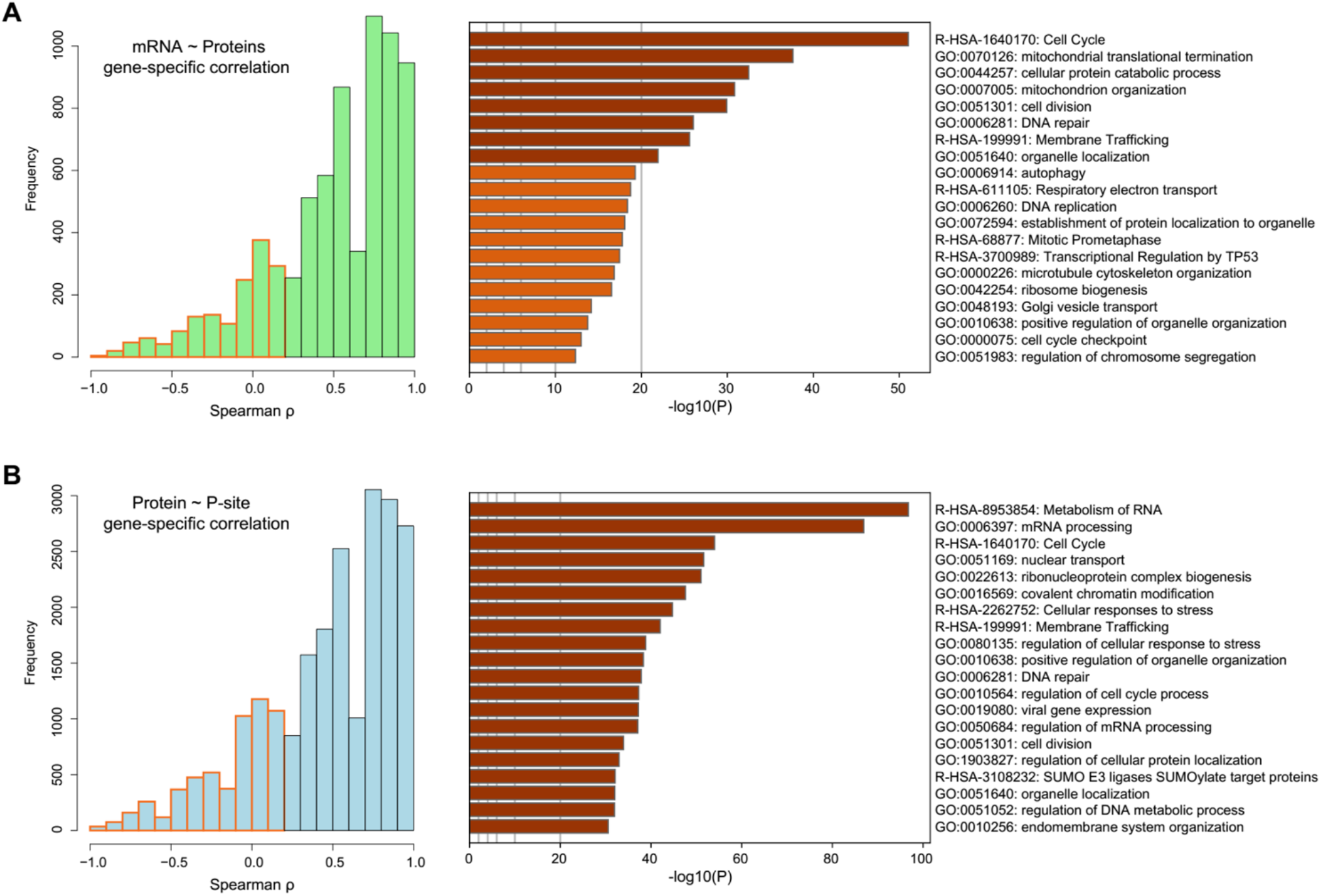
The identification and functional annotation of the genes with low correlations across cells. (A) The histogram of the within-gene correlations between mRNA and protein. The genes with low correlation (ρ < 0.2, Spearman correlation) were marked in orange and functionally annotated by Meatscape. (B) The histogram of the within-phosphosite correlations between protein and phosphosite. The genes with low correlations (ρ < 0.2) were marked in red and functionally annotated by Meatscape.

In summary, the across-gene mRNA~protein and Protein~P-site correlations could be determined at both absolute and relative scales using our high-quality DIA-MS datasets. Together with the biological annotations of gene-specific correlations, our integrative analysis uncovered ubiquitous but also varying determinants for protein expression and phosphorylation levels. The relative-scale analyses indicate that mRNA levels cannot fully predict protein abundance and that protein abundance cannot fully predict the phosphorylation level variability. For the genes in specific pathways, the protein and phosphoprotein-level alterations may be even more difficult to predict than others. Thus, the accurate proteotyping of a cell line favors real experimental data.

### 3.5 Comparing different phosphoproteomic normalization strategies using the matched proteome and phosphoproteome DIA-MS datasets

Many of the previous phosphoproteomic studies only measured the relative changes between samples based on the enriched phosphopeptides. However, as shown by the relative Protein~P-site correlation between stable cancer cells (**Figure 4**) and similar analysis performed in the previous studies ^51-53^, a major component underlying P-site abundance variance is attributed to the protein levels. It is conceivable that this complication also occurs in other steady-state comparison scenarios, such as comparing the phosphoproteomes of the tumor and paired adjacent tissues. Although many recent clinical proteomic studies measured both total proteomes and phosphoproteomes across samples, these two layered datasets are usually quantified respectively. Wu *et al.* showed that 25% of the phosphopeptide abundance changes could be due to protein expression differences in a mutant yeast system, emphasizing the critical importance of calibrating phosphoproteomic data by protein expression ^52^. On the other hand, the protein-level measurement is not straightforward in bottom-up proteomics, in which peptide abundances are measured as surrogates for protein expressions ^53^. Currently, there is no consensus on how to summarize peptide’s abundance into protein levels; and a common procedure is to summarize signals of Top-intensity peptides for a particular protein (*e.g.*, the “Top 1” or “Top 3” method). However, if Top-intensity peptides are used for normalizing phosphoproteomic data, one has to assume that e.g., these peptides are not modified themselves, or their modifications would not affect the relative protein quantification (**Supplementary Figure S3A**). Herein, based on the matched peptide and phosphopeptide DIA datasets, we investigate the influence of different normalization strategies on phosphoproteomic profiling.

To acquire the site-specific, quantitative phosphoproteomic data, we adopted three possible methods (**Figure 6A**) - **Method 1**: the intensity of phosphopeptides (P-peptide) is used directly without protein-level information; **Method 2**: the intensity of P-peptides is normalized so that it is divided by the protein expression that is estimated by Top 3 method ^27, 54^ in the total proteomic measurement; and **Method 3**: the intensity of P-peptides is normalized so that it is divided by the abundance of *its sequence-matched, unmodified peptide counterpart* (***nP-peptide***) in the total proteomic measurement. We included Method 3 because we reason that the ratio of *<u>P-peptide/nP-peptide</u>* is independent of the modification status of all the other peptides of the same protein and may be more robust for phosphosite abundance normalization. As expected, because Method 3 requires the identification of both P- and nP-peptides, it only determined the quantification ratio for 9,271 phosphosites, which accounts for 37.9% of those analyzed by Method 1 (**Figure 6B**). In contrast, Method 2 retains the quantitative information for 90.3% of phosphopeptides that were analyzed by Method 1. The correlation analysis across phosphosites indicated that whereas Method 2 and 3 were highly correlated among cells (R was around 0.8), Method 1 generated deviated quantification results to Method 2 and 3 (R was around 0.6, **Figure 6C**). The HCA after scaling also suggested that Method 2 and 3 results were clustered together (**Supplementary Figure S3B)**. Furthermore, Method 1 and 2 respectively identified 574 and 550 phosphosites changing significantly between the metastatic and primary melanoma group, with 145 of them overlapping between Method 1 and Method 2 (**Figure 6D**). Therefore, our results showed that the different phosphoproteomic normalization strategies had a significant effect on both absolute and relative phosphosite quantification. Finally, we correlated the *P-peptide/Top3-peptide* (ratio in Method 2) and *P-peptide/nP-peptide* (ratio in Method 3) across the six cell lines for each phosphosite. We found that the relative change of ratios was largely consistent between Method 2 and 3 (R=0.817, averaged from 9,217 phosphosites, **Supplementary Figure S3C**). Also, the relative fold-change of Metastatic vs. Primary cells was generally conserved between results following Method 2 or 3 (R=0.61, **Figure 6E**). Given the fact that Method 2 analyzed 140% more phosphosites than Method 3, we conclude that Method 2 efficiently accounted for the protein expression difference in quantifying phosphosite while still maintaining a fairly large phosphoproteome coverage. Therefore, Method 2 could be accepted as the initial normalization strategy for large-scale phosphoproteomic analysis.

**Figure 6.**
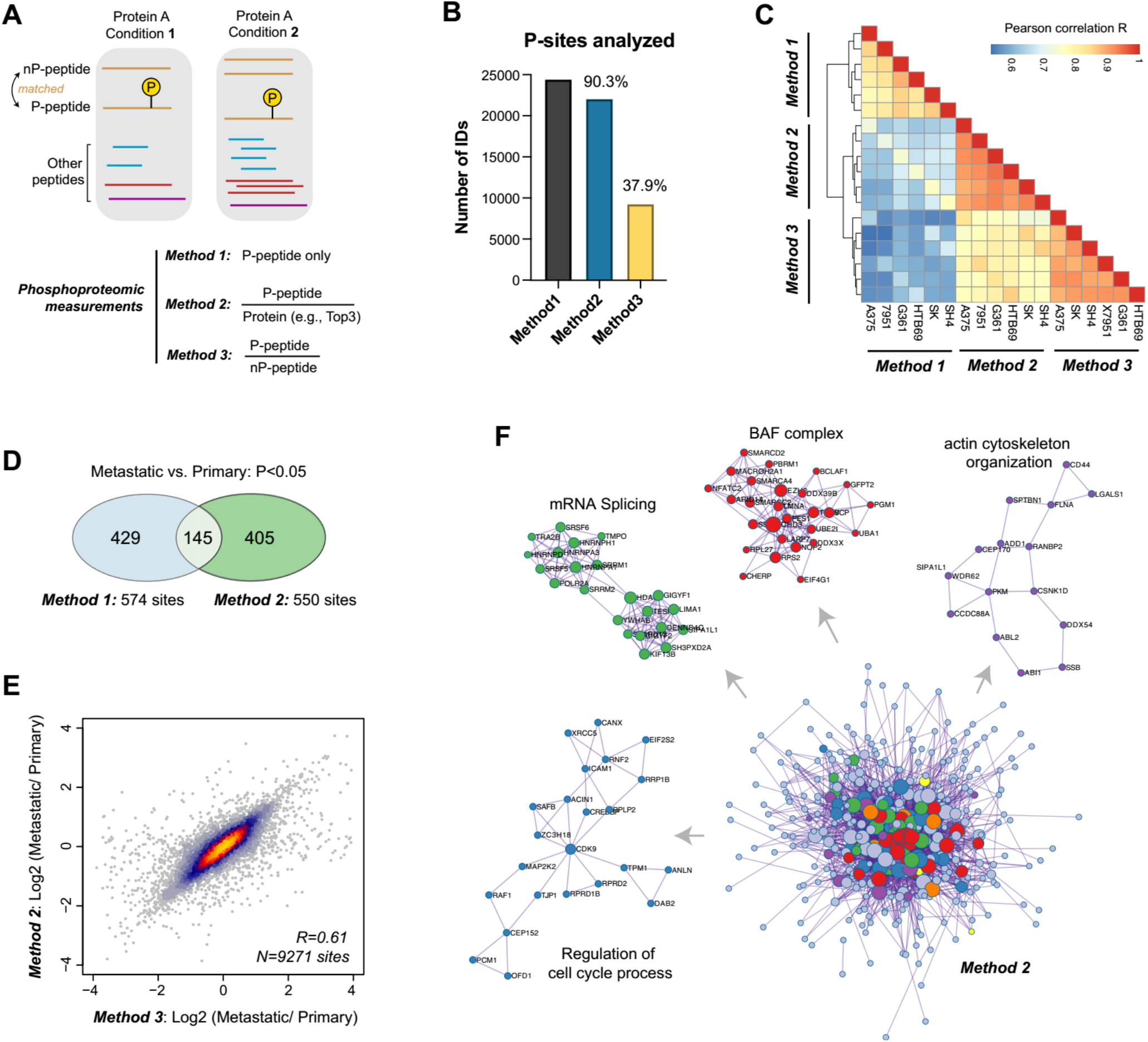
The comparisons between different phosphoproteomic normalization strategies and phosphorylation-centric signaling alterations between metastatic and non-metastatic melanoma cell lines. (A) The overall scheme of three phosphoproteomic normalization strategies. (B) The number of P-sites analyzed by three methods. (C) The correlation analysis of P-sites analyzed by three methods. (D) The overlapped P-sites that were significantly changed between Method 1 and Method 2. (E) The scatter plot of the log-scale fold-changes of 9,271 P-sites between three metastatic cells and three primary cells determined by Method 2 or Method 3. (F) Following Method 2, 402 metastasis-associated genes were filtered (*P*<0.05 between metastatic and primary groups, student’s t-test) for functional annotation and the protein-protein interaction (PPI) network analysis by Metascape. The MCODE components were identified from the merged PPI network as core phosphoprotein modules and highlighted in different colors.

### 3.6 Preliminary biological insights on melanoma cancer metastasis using multi-omic profiling

Because of the small sample size and the usage of cell lines (rather than clinical samples), this study did not aim to identify biomarkers associated with cancer metastasis in melanoma. Nevertheless, we discovered that the individual genomic variability, rather than the metastatic property, determined the overall molecular similarity between six cell lines (**Figure 2E-H**). In fact, simple t-tests filtered 311, 242, and 425 genes being different at mRNA, proteomic, and phosphoproteomic levels, respectively (*P*<0.05 as a loose criterion), accounting for only 2.30%, 2.87%, and 6.62% of the total genes measured at each level (see the numbers in **Figure 2B**). We therefore included all these differential genes for function analysis, with the purpose to discern metastasis-related processes at each level. According to Metascape ^37^, we found that different omics layers uncovered many distinctive signature genes associated with metastasis, with a few overlapping functional processes across layers, such as DNA repair, Metabolism of mRNA, Negative regulation of cell cycle, and Regulation of intracellular transport (*P* <0.01 for all, **Supplementary Figure S4A-B**). We further found that the phosphoproteomic dataset particularly revealed more metastasis-associated signaling processes than proteomic profiling, such as SWI/SNF complex, the beta-catenin-TCF complex assembly, and mRNA processing (all *P* <0.01). Intriguingly, the proteome data uniquely revealed that Glycosylphosphatidylinositol (GPI)-anchor transamidase complex was significantly changed between Metastatic and Primary cells (-Log10_P value =10.33, **Supplementary Figure S4C-D**). In particular, all the five subunits of this complex-GPAA1, PIGK, PIGT, PIGS, PIGU were found to upregulated by approximately 2 folds in the three metastatic cell lines, compared to the three primary cells (**Supplementary Figure S5A**). The inspection of the published mRNA profiles through the Online consensus Survival webserver for Skin Cutaneous Melanoma (OSskcm)^38^ supported the unfavorable prognosis outcome for GPAA1 (*P*=5E-04 for 1085 clinical melanoma samples combined, *P*=0.022 for TCGA datasets, **Supplementary Figure S5B**). Moreover, previous literature has suggested that the overexpression of GPI transamidase subunits induces tumor invasion in breast cancer ^55^ and that GPAA1 promotes gastric cancer progression via upregulation of GPI-anchored proteins ^56^. It is therefore appealing to establish the functional relationship between GPI transamidase complex overexpression and melanoma metastasis in the future biological and clinical studies.

Finally, to focus on phosphoproteomic data that enrich cell signaling events, we normalized the phosphosite abundance by the corresponding proteins (following *Method 2 in Section 3.5*) and took the 402 filtered genes (*P*<0.05, student’s t-test) for functional analysis. Here, all protein-protein interactions (PPI) among the 402 gene list were extracted from PPI data source in Metascape ^37^, and core protein modules were then identified in the PPI network by MCODE ^36^ (**Figure 6F**). These protein modules include BAF complex, actin cytoskeleton organization, mRNA splicing, and Regulation of cell cycle processes, providing an overview of those most important phosphorylation-centric signaling alterations between metastatic and non-metastatic melanoma cell lines tested. The establishment of relevant molecular mechanisms connecting the phosphoprotein module to melanoma development is beyond the scope of this study.

## 4 Conclusions

In this study, we presented a high-quality DIA-MS dataset, which profiled both proteome and phosphoproteome in the six melanoma cell lines that differ in the metastatic cancer phenotype. With the respective 2-hour single-shot measurements, DIA-MS achieved a proteomic coverage of 8,100 proteins and a phosphoproteomic coverage of 40,000 phosphopeptides for each cancer cell line, demonstrating the exquisite sensitivity and sample throughput of DIA-MS in cell proteotyping. Besides the substantial genome-wide analyte throughput, DIA-MS showed an equally excellent quantification reproducibility compared to RNA-seq. The additional comparison of our DIA-MS results to CCLE-TMT data also revealed comparable qualitative and quantitative features between the two proteomic methods. Considering its flexibility of handling individual cell lines, the cost-effectiveness, the simple experimental procedures, and the analytical performance newly available from the spectral-library free workflow, as well as the recently demonstrated cross-lab reproducibility ^27, 57^, we deem DIA-MS a powerful and competitive method for documenting and referencing the gene expression landscape of cancer cells used in a particular experiment and between labs ^10^.

Human cancer cells are widely used for large-scale comparative studies across multi-omic layers. Benefiting from the high data-quality of DIA-MS and the well-matched multi-omic datasets generated in the present study, we quantitatively assessed the across-gene mRNA~protein correlation and Protein~P-site correlation at both absolute and relative scales. We then summarized the biological processes associated with the low gene-specific correlations across cell lines. We additionally benchmarked the possible phosphoproteomic normalization strategies. We finally searched our multi-omic datasets for potential biological signatures and insights separating metastatic and primary melanoma cells. Our analysis demonstrated the significant dependence of protein abundances on mRNA concentrations (ρ= 0.61) and the significant dependence of phosphosite regulation on protein changes (ρ= 0.53). Furthermore, our data supported the fundamental need of calibrating phosphoproteomic abundance by corresponding protein expression and suggested that the Top-intensity method could be acceptable to summarize protein intensities for this normalization purpose. Finally, our results underscored that distinctive metastatic signatures could emerge at different molecular layers and that both proteomic and phosphoproteomic measurements are indispensable for the complete understanding of biological processes.

## Acknowledgements

Y.L. thanks the support from the National Institute of General Medical Sciences (NIGMS), National Institutes of Health (NIH) through Grant R01GM137031 to Y.L. This research was supported in part by a Pilot Grant from Yale Cancer Center.

## Notes

The authors declare no competing interests.

## Author Contributions

E.G. and C.W. performed cell-culturing and conducted the transcriptomic, proteomic and phosphoproteomic sample procession experiments for all cancer cell lines. W.L. performed MS measurements. W.S. and Y.D. provided critical comments to the study and manuscript. Y.L. wrote the paper based on the inputs from all the authors. Y.L. supervised the study.

## References

1. T. Geiger, A. Wehner, C. Schaab, J. Cox and M. Mann, Molecular & cellular proteomics: MCP, 2012, 11, M111 014050.

2. J. Barretina, G. Caponigro, N. Stransky, K. Venkatesan, A. A. Margolin, S. Kim, C. J. Wilson, J. Lehar, G. V. Kryukov, D. Sonkin, A. Reddy, M. Liu, L. Murray, M. F. Berger, J. E. Monahan, P. Morais, J. Meltzer, A. Korejwa, J. Jane-Valbuena, F. A. Mapa, J. Thibault, E. Bric-Furlong, P. Raman, A. Shipway, I. H. Engels, J. Cheng, G. K. Yu, J. Yu, P. Aspesi, Jr., M. de Silva, K. Jagtap, M. D. Jones, L. Wang, C. Hatton, E. Palescandolo, S. Gupta, S. Mahan, C. Sougnez, R. C. Onofrio, T. Liefeld, L. MacConaill, W. Winckler, M. Reich, N. Li, J. P. Mesirov, S. B. Gabriel, G. Getz, K. Ardlie, V. Chan, V. E. Myer, B. L. Weber, J. Porter, M. Warmuth, P. Finan, J. L. Harris, M. Meyerson, T. R. Golub, M. P. Morrissey, W. R. Sellers, R. Schlegel and L. A. Garraway, Nature, 2012, 483, 603–607.

3. M. Ghandi, F. W. Huang, J. Jane-Valbuena, G. V. Kryukov, C. C. Lo, E. R. McDonald, 3rd, J. Barretina, E. T. Gelfand, C. M. Bielski, H. Li, K. Hu, A. Y. Andreev-Drakhlin, J. Kim, J. M. Hess, B. J. Haas, F. Aguet, B. A. Weir, M. V. Rothberg, B. R. Paolella, M. S. Lawrence, R. Akbani, Y. Lu, H. L. Tiv, P. C. Gokhale, A. de Weck, A. A. Mansour, C. Oh, J. Shih, K. Hadi, Y. Rosen, J. Bistline, K. Venkatesan, A. Reddy, D. Sonkin, M. Liu, J. Lehar, J. M. Korn, D. A. Porter, M. D. Jones, J. Golji, G. Caponigro, J. E. Taylor, C. M. Dunning, A. L. Creech, A. C. Warren, J. M. McFarland, M. Zamanighomi, A. Kauffmann, N. Stransky, M. Imielinski, Y. E. Maruvka, A. D. Cherniack, A. Tsherniak, F. Vazquez, J. D. Jaffe, A. A. Lane, D. M. Weinstock, C. M. Johannessen, M. P. Morrissey, F. Stegmeier, R. Schlegel, W. C. Hahn, G. Getz, G. B. Mills, J. S. Boehm, T. R. Golub, L. A. Garraway and W. R. Sellers, Nature, 2019, 569, 503–508.

4. D. P. Nusinow, J. Szpyt, M. Ghandi, C. M. Rose, E. R. McDonald, 3rd, M. Kalocsay, J. Jane-Valbuena, E. Gelfand, D. K. Schweppe, M. Jedrychowski, J. Golji, D. A. Porter, T. Rejtar, Y. K. Wang, G. V. Kryukov, F. Stegmeier, B. K. Erickson, L. A. Garraway, W. R. Sellers and S. P. Gygi, Cell, 2020, 180, 387–402 e316.

5. T. I. Roumeliotis, S. P. Williams, E. Goncalves, C. Alsinet, M. Del Castillo Velasco-Herrera, N. Aben, F. Z. Ghavidel, M. Michaut, M. Schubert, S. Price, J. C. Wright, L. Yu, M. Yang, R. Dienstmann, J. Guinney, P. Beltrao, A. Brazma, M. Pardo, O. Stegle, D. J. Adams, L. Wessels, J. Saez-Rodriguez, U. McDermott and J. S. Choudhary, Cell reports, 2017, 20, 2201–2214.

6. T. Guo, A. Luna, V. N. Rajapakse, C. C. Koh, Z. Wu, W. Liu, Y. Sun, H. Gao, M. P. Menden, C. Xu, L. Calzone, L. Martignetti, C. Auwerx, M. Buljan, A. Banaei-Esfahani, A. Ori, M. Iskar, L. Gillet, R. Bi, J. Zhang, H. Zhang, C. Yu, Q. Zhong, S. Varma, U. Schmitt, P. Qiu, Q. Zhang, Y. Zhu, P. J. Wild, M. J. Garnett, P. Bork, M. Beck, K. Liu, J. Saez-Rodriguez, F. Elloumi, W. C. Reinhold, C. Sander, Y. Pommier and R. Aebersold, iScience, 2019, 21, 664–680.

7. A. M. Gholami, H. Hahne, Z. Wu, F. J. Auer, C. Meng, M. Wilhelm and B. Kuster, Cell reports, 2013, 4, 609–620.

8. M. Frejno, C. Meng, B. Ruprecht, T. Oellerich, S. Scheich, K. Kleigrewe, E. Drecoll, P. Samaras, A. Hogrebe, D. Helm, J. Mergner, J. Zecha, S. Heinzlmeir, M. Wilhelm, J. Dorn, H. M. Kvasnicka, H. Serve, W. Weichert and B. Kuster, Nature communications, 2020, 11, 3639.

9. U. Ben-David, B. Siranosian, G. Ha, H. Tang, Y. Oren, K. Hinohara, C. A. Strathdee, J. Dempster, N. J. Lyons, R. Burns, A. Nag, G. Kugener, B. Cimini, P. Tsvetkov, Y. E. Maruvka, R. O’Rourke, A. Garrity, A. A. Tubelli, P. Bandopadhayay, A. Tsherniak, F. Vazquez, B. Wong, C. Birger, M. Ghandi, A. R. Thorner, J. A. Bittker, M. Meyerson, G. Getz, R. Beroukhim and T. R. Golub, Nature, 2018, 560, 325–330.

10. Y. Liu, Y. Mi, T. Mueller, S. Kreibich, E. G. Williams, A. Van Drogen, C. Borel, M. Frank, P. L. Germain, I. Bludau, M. Mehnert, M. Seifert, M. Emmenlauer, I. Sorg, F. Bezrukov, F. S. Bena, H. Zhou, C. Dehio, G. Testa, J. Saez-Rodriguez, S. E. Antonarakis, W. D. Hardt and R. Aebersold, Nature biotechnology, 2019, 37, 314–322.

11. M. Yu, S. K. Selvaraj, M. M. Liang-Chu, S. Aghajani, M. Busse, J. Yuan, G. Lee, F. Peale, C. Klijn, R. Bourgon, J. S. Kaminker and R. M. Neve, Nature, 2015, 520, 307–311.

12. P. Picotti, M. Clement-Ziza, H. Lam, D. S. Campbell, A. Schmidt, E. W. Deutsch, H. Rost, Z. Sun, O. Rinner, L. Reiter, Q. Shen, J. J. Michaelson, A. Frei, S. Alberti, U. Kusebauch, B. Wollscheid, R. L. Moritz, A. Beyer and R. Aebersold, Nature, 2013, 494, 266–270.

13. Y. Jiang, A. Sun, Y. Zhao, W. Ying, H. Sun, X. Yang, B. Xing, W. Sun, L. Ren, B. Hu, C. Li, L. Zhang, G. Qin, M. Zhang, N. Chen, M. Zhang, Y. Huang, J. Zhou, Y. Zhao, M. Liu, X. Zhu, Y. Qiu, Y. Sun, C. Huang, M. Yan, M. Wang, W. Liu, F. Tian, H. Xu, J. Zhou, Z. Wu, T. Shi, W. Zhu, J. Qin, L. Xie, J. Fan, X. Qian, F. He and C. Chinese Human Proteome Project, Nature, 2019, 567, 257–261.

14. P. Mertins, D. R. Mani, K. V. Ruggles, M. A. Gillette, K. R. Clauser, P. Wang, X. Wang, J. W. Qiao, S. Cao, F. Petralia, E. Kawaler, F. Mundt, K. Krug, Z. Tu, J. T. Lei, M. L. Gatza, M. Wilkerson, C. M. Perou, V. Yellapantula, K. L. Huang, C. Lin, M. D. McLellan, P. Yan, S. R. Davies, R. R. Townsend, S. J. Skates, J. Wang, B. Zhang, C. R. Kinsinger, M. Mesri, H. Rodriguez, L. Ding, A. G. Paulovich, D. Fenyo, M. J. Ellis, S. A. Carr and C. Nci, Nature, 2016, 534, 55–62.

15. Q. Gao, H. Zhu, L. Dong, W. Shi, R. Chen, Z. Song, C. Huang, J. Li, X. Dong, Y. Zhou, Q. Liu, L. Ma, X. Wang, J. Zhou, Y. Liu, E. Boja, A. I. Robles, W. Ma, P. Wang, Y. Li, L. Ding, B. Wen, B. Zhang, H. Rodriguez, D. Gao, H. Zhou and J. Fan, Cell, 2019, 179, 561–577 e522.

16. D. J. Clark, S. M. Dhanasekaran, F. Petralia, J. Pan, X. Song, Y. Hu, F. da Veiga Leprevost, B. Reva, T. M. Lih, H. Y. Chang, W. Ma, C. Huang, C. J. Ricketts, L. Chen, A. Krek, Y. Li, D. Rykunov, Q. K. Li, L. S. Chen, U. Ozbek, S. Vasaikar, Y. Wu, S. Yoo, S. Chowdhury, M. A. Wyczalkowski, J. Ji, M. Schnaubelt, A. Kong, S. Sethuraman, D. M. Avtonomov, M. Ao, A. Colaprico, S. Cao, K. C. Cho, S. Kalayci, S. Ma, W. Liu, K. Ruggles, A. Calinawan, Z. H. Gumus, D. Geiszler, E. Kawaler, G. C. Teo, B. Wen, Y. Zhang, S. Keegan, K. Li, F. Chen, N. Edwards, P. M. Pierorazio, X. S. Chen, C. P. Pavlovich, A. A. Hakimi, G. Brominski, J. J. Hsieh, A. Antczak, T. Omelchenko, J. Lubinski, M. Wiznerowicz, W. M. Linehan, C. R. Kinsinger, M. Thiagarajan, E. S. Boja, M. Mesri, T. Hiltke, A. I. Robles, H. Rodriguez, J. Qian, D. Fenyo, B. Zhang, L. Ding, E. Schadt, A. M. Chinnaiyan, Z. Zhang, G. S. Omenn, M. Cieslik, D. W. Chan, A. I. Nesvizhskii, P. Wang, H. Zhang and C. Clinical Proteomic Tumor Analysis, Cell, 2019, 179, 964–983 e931.

17. L. C. Gillet, P. Navarro, S. Tate, H. Rost, N. Selevsek, L. Reiter, R. Bonner and R. Aebersold, Molecular & cellular proteomics: MCP, 2012, 11, O111 016717.

18. R. Aebersold and M. Mann, Nature, 2016, 537, 347–355.

19. J. D. Venable, M. Q. Dong, J. Wohlschlegel, A. Dillin and J. R. Yates, Nature methods, 2004, 1, 39–45.

20. C. Li, Y. D. Sun, G. Y. Yu, J. R. Cui, Z. Lou, H. Zhang, Y. Huang, C. G. Bai, L. L. Deng, P. Liu, K. Zheng, Y. H. Wang, Q. Q. Wang, Q. R. Li, Q. Q. Wu, Q. Liu, Y. Shyr, Y. X. Li, L. N. Chen, J. R. Wu, W. Zhang and R. Zeng, Cancer Cell, 2020, 38, 734–747 e739.

21. J. Muntel, J. Kirkpatrick, R. Bruderer, T. Huang, O. Vitek, A. Ori and L. Reiter, Journal of proteome research, 2019, 18, 1340–1351.

22. C. Everaert, M. Luypaert, J. L. V. Maag, Q. X. Cheng, M. E. Dinger, J. Hellemans and P. Mestdagh, Sci Rep, 2017, 7, 1559.

23. D. Kim, J. M. Paggi, C. Park, C. Bennett and S. L. Salzberg, Nature biotechnology, 2019, 37, 907–915.

24. A. Frankish, M. Diekhans, A. M. Ferreira, R. Johnson, I. Jungreis, J. Loveland, J. M. Mudge, C. Sisu, J. Wright, J. Armstrong, I. Barnes, A. Berry, A. Bignell, S. Carbonell Sala, J. Chrast, F. Cunningham, T. Di Domenico, S. Donaldson, I. T. Fiddes, C. Garcia Giron, J. M. Gonzalez, T. Grego, M. Hardy, T. Hourlier, T. Hunt, O. G. Izuogu, J. Lagarde, F. J. Martin, L. Martinez, S. Mohanan, P. Muir, F. C. P. Navarro, A. Parker, B. Pei, F. Pozo, M. Ruffier, B. M. Schmitt, E. Stapleton, M. M. Suner, I. Sycheva, B. Uszczynska-Ratajczak, J. Xu, A. Yates, D. Zerbino, Y. Zhang, B. Aken, J. S. Choudhary, M. Gerstein, R. Guigo, T. J. P. Hubbard, M. Kellis, B. Paten, A. Reymond, M. L. Tress and P. Flicek, Nucleic Acids Res, 2019, 47, D766–D773.

25. M. Pertea, D. Kim, G. M. Pertea, J. T. Leek and S. L. Salzberg, Nat Protoc, 2016, 11, 1650–1667.

26. W. Li, H. Chi, B. Salovska, C. Wu, L. Sun, G. Rosenberger and Y. Liu, J Am Soc Mass Spectrom, 2019, DOI: 10.1007/s13361-019-02243-1.

27. B. C. Collins, C. L. Hunter, Y. Liu, B. Schilling, G. Rosenberger, S. L. Bader, D. W. Chan, B. W. Gibson, A. C. Gingras, J. M. Held, M. Hirayama-Kurogi, G. Hou, C. Krisp, B. Larsen, L. Lin, S. Liu, M. P. Molloy, R. L. Moritz, S. Ohtsuki, R. Schlapbach, N. Selevsek, S. N. Thomas, S. C. Tzeng, H. Zhang and R. Aebersold, Nature communications, 2017, 8, 291.

28. M. Mehnert, W. Li, C. Wu, B. Salovska and Y. Liu, Proteomics, 2019, DOI: 10.1002/pmic.201800438, e1800438.

29. R. Bruderer, O. M. Bernhardt, T. Gandhi, S. M. Miladinovic, L. Y. Cheng, S. Messner, T. Ehrenberger, V. Zanotelli, Y. Butscheid, C. Escher, O. Vitek, O. Rinner and L. Reiter, Molecular & cellular proteomics: MCP, 2015, 14, 1400–1410.

30. R. Bruderer, O. M. Bernhardt, T. Gandhi, Y. Xuan, J. Sondermann, M. Schmidt, D. Gomez-Varela and L. Reiter, Molecular & cellular proteomics: MCP, 2017, 16, 2296–2309.

31. C. C. Tsou, D. Avtonomov, B. Larsen, M. Tucholska, H. Choi, A. C. Gingras and A. I. Nesvizhskii, Nature methods, 2015, 12, 258–264, 257 p following 264.

32. G. Rosenberger, Y. Liu, H. L. Rost, C. Ludwig, A. Buil, A. Bensimon, M. Soste, T. D. Spector, E. T. Dermitzakis, B. C. Collins, L. Malmstrom and R. Aebersold, Nature biotechnology, 2017, 35, 781–788.

33. D. B. Bekker-Jensen, O. M. Bernhardt, A. Hogrebe, A. Martinez-Val, L. Verbeke, T. Gandhi, C. D. Kelstrup, L. Reiter and J. V. Olsen, Nature communications, 2020, 11, 787.

34. J. V. Olsen, B. Blagoev, F. Gnad, B. Macek, C. Kumar, P. Mortensen and M. Mann, Cell, 2006, 127, 635–648.

35. P. Shannon, A. Markiel, O. Ozier, N. S. Baliga, J. T. Wang, D. Ramage, N. Amin, B. Schwikowski and T. Ideker, Genome research, 2003, 13, 2498–2504.

36. G. D. Bader and C. W. Hogue, BMC bioinformatics, 2003, 4, 2.

37. Y. Zhou, B. Zhou, L. Pache, M. Chang, A. H. Khodabakhshi, O. Tanaseichuk, C. Benner and S. K. Chanda, Nature communications, 2019, 10, 1523.

38. L. Zhang, Q. Wang, L. Wang, L. Xie, Y. An, G. Zhang, W. Zhu, Y. Li, Z. Liu, X. Zhang, P. Tang, X. Huo and X. Guo, Cancer Cell Int, 2020, 20, 176.

39. Y. Perez-Riverol, A. Csordas, J. Bai, M. Bernal-Llinares, S. Hewapathirana, D. J. Kundu, A. Inuganti, J. Griss, G. Mayer, M. Eisenacher, E. Perez, J. Uszkoreit, J. Pfeuffer, T. Sachsenberg, S. Yilmaz, S. Tiwary, J. Cox, E. Audain, M. Walzer, A. F. Jarnuczak, T. Ternent, A. Brazma and J. A. Vizcaino, Nucleic Acids Res, 2019, 47, D442–D450.

40. M. Beck, A. Schmidt, J. Malmstroem, M. Claassen, A. Ori, A. Szymborska, F. Herzog, O. Rinner, J. Ellenberg and R. Aebersold, Molecular systems biology, 2011, 7, 549.

41. Y. Liu, A. Beyer and R. Aebersold, Cell, 2016, 165, 535–550.

42. C. Buccitelli and M. Selbach, Nature reviews. Genetics, 2020, 21, 630–644.

43. G. C. McAlister, D. P. Nusinow, M. P. Jedrychowski, M. Wuhr, E.L. Huttlin, B. K. Erickson, R. Rad, W. Haas and S. P. Gygi, Analytical chemistry, 2014, 86, 7150–7158.

44. X. Liu, S. P. Gygi and J. A. Paulo, Proteomics, 2020, DOI: 10.1002/pmic.202000218, e2000218.

45. O. T. Schubert, C. Ludwig, M. Kogadeeva, M. Zimmermann, G. Rosenberger, M. Gengenbacher, L. C. Gillet, B. C. Collins, H. L. Rost, S. H. Kaufmann, U. Sauer and R. Aebersold, Cell Host Microbe, 2015, 18, 96–108.

46. N. Fortelny, C. M. Overall, P. Pavlidis and G. V. C. Freue, Nature, 2017, 547, E19–E20.

47. G. M. Silva and C. Vogel, Mol Syst Biol, 2016, 12, 885.

48. B. Salovska, H. Zhu, T. Gandhi, M. Frank, W. Li, G. Rosenberger, C. Wu, P. L. Germain, H. Zhou, Z. Hodny, L. Reiter and Y. Liu, Molecular systems biology, 2020, 16, e9170.

49. J. Zecha, C. Meng, D. P. Zolg, P. Samaras, M. Wilhelm and B. Kuster, Mol Cell Proteomics, 2018, 17, 974–992.

50. H. Zhang, T. Liu, Z. Zhang, S. H. Payne, B. Zhang, J. E. McDermott, J. Y. Zhou, V. A. Petyuk, L. Chen, D. Ray, S. Sun, F. Yang, L. Chen, J. Wang, P. Shah, S. W. Cha, P. Aiyetan, S. Woo, Y. Tian, M. A. Gritsenko, T. R. Clauss, C. Choi, M. E. Monroe, S. Thomas, S. Nie, C. Wu, R. J. Moore, K. H. Yu, D. L. Tabb, D. Fenyo, V. Bafna, Y. Wang, H. Rodriguez, E. S. Boja, T. Hiltke, R. C. Rivers, L. Sokoll, H. Zhu, I. M. Shih, L. Cope, A. Pandey, B. Zhang, M. P. Snyder, D. A. Levine, R. D. Smith, D. W. Chan, K. D. Rodland and C. Investigators, Cell, 2016, 166, 755–765.

51. Y. B. Wu, J. Dai, X. L. Yang, S. J. Li, S. L. Zhao, Q. H. Sheng, J. S. Tang, G. Y. Zheng, Y. X. Li, J. R. Wu and R. Zeng, Molecular & cellular proteomics: MCP, 2009, 8, 2809–2826.

52. R. Wu, N. Dephoure, W. Haas, E. L. Huttlin, B. Zhai, M. E. Sowa and S. P. Gygi, Molecular & cellular proteomics: MCP, 2011, 10, M111 009654.

53. J. V. Olsen, M. Vermeulen, A. Santamaria, C. Kumar, M. L. Miller, L. J. Jensen, F. Gnad, J. Cox, T. S. Jensen, E. A. Nigg, S. Brunak and M. Mann, Sci Signal, 2010, 3, ra3.

54. G. Rosenberger, C. Ludwig, H. L. Rost, R. Aebersold and L. Malmstrom, Bioinformatics, 2014, 30, 2511–2513.

55. G. Wu, Z. Guo, A. Chatterjee, X. Huang, E. Rubin, F. Wu, E. Mambo, X. Chang, M. Osada, M. Sook Kim, C. Moon, J. A. Califano, E. A. Ratovitski, S. M. Gollin, S. Sukumar, D. Sidransky and B. Trink, Cancer Res, 2006, 66, 9829–9836.

56. X. X. Zhang, B. Ni, Q. Li, L. P. Hu, S. H. Jiang, R. K. Li, G. A. Tian, L. L. Zhu, J. Li, X. L. Zhang, Y. L. Zhang, X. M. Yang, Q. Yang, Y. H. Wang, C. C. Zhu and Z. G. Zhang, J Exp Clin Cancer Res, 2019, 38, 214.

57. Y. Xuan, N. W. Bateman, S. Gallien, S. Goetze, Y. Zhou, P. Navarro, M. Hu, N. Parikh, B. L. Hood, K. A. Conrads, C. Loosse, R. B. Kitata, S. R. Piersma, D. Chiasserini, H. Zhu, G. Hou, M. Tahir, A. Macklin, A. Khoo, X. Sun, B. Crossett, A. Sickmann, Y. J. Chen, C. R. Jimenez, H. Zhou, S. Liu, M. R. Larsen, T. Kislinger, Z. Chen, B. L. Parker, S. J. Cordwell, B. Wollscheid and T. P. Conrads, Nature communications, 2020, 11, 5248.

