## Supplementary Figures S1-S5 for "Data-independent Acquisition-based Proteome and Phosphoproteome Profiling across Six Melanoma Cell Lines Reveals Determinants of Proteotypes"

**Supporting Figures**

**Supporting Figures** ........ 2

*Supplementary Figure S1* ........ 2

*Supplementary Figure S2* ........ 3

*Supplementary Figure S3* ........ 4

*Supplementary Figure S4* ........ 5

*Supplementary Figure S5* ........ 6

**Supplementary Table**

***Supplementary Table S1 | The quantitative results at mRNA, protein, and phosphosite levels across the six melanoma cell lines.*** (Table will be available upon paper publication).

### Supporting Figures (S1-S5)

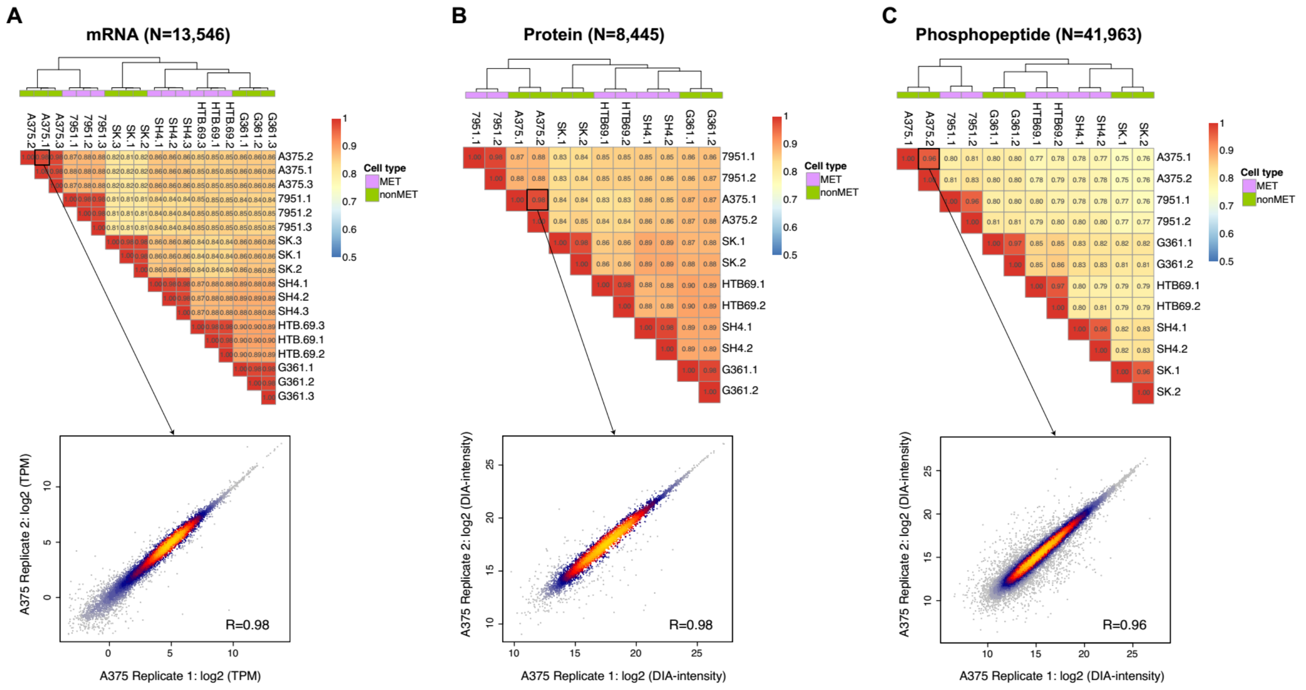


**Supplementary Figure S1 | The correlation analysis of TPM values or DIA-MS peak areas between dish-replicates for each line, grouped by mRNA (A), protein (B) and phosphopeptide (C).** Dish-replicates were clustered according to the datasets of all three molecular layers, with equally excellent quantification reproducibility.

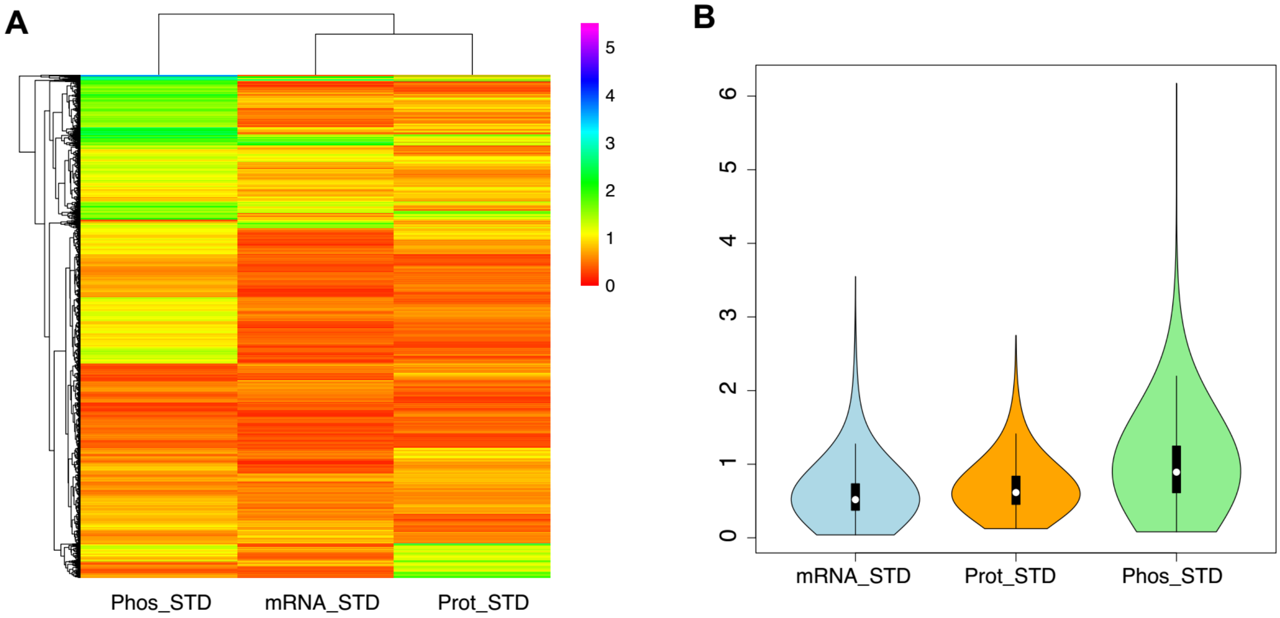


**Supplementary Figure S2 | The quantitative viability, measured as standard deviation (STD) across the six cell lines, for mRNA, protein and phosphopeptide, shown as (A) the heatmap and (B) the violin plot.**

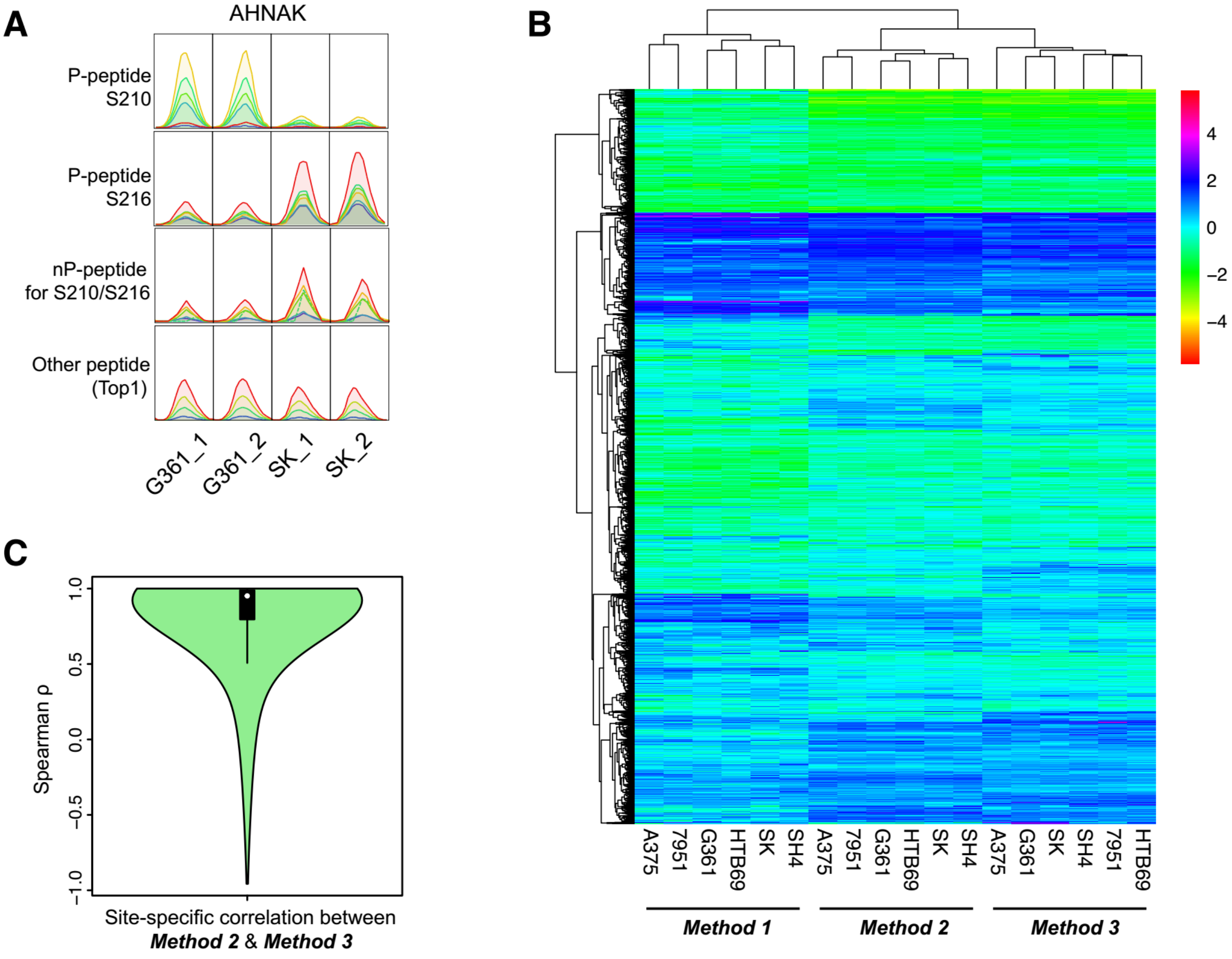


**Supplementary Figure S3 | Application and analysis of different phosphoproteomic normalization strategies.** (A) A real-world example of the intensities of a phosphopeptide and its associated counterparts. The S210 and S216 of Neuroblast differentiation-associated protein AHNAK protein had distinctive quantification ratios between G361 and SK cells. The nP-peptide of both phosphosites was determined to follow the quantitative ratio of S216 but not S210, whereas the Top 1 peptide only presented a fold-change much smaller than that of the nP-peptide. In this particular case, S216 would be regarded as non-changed phosphosite between cells, if nP-peptide was used as reference. (B) The HCA analysis of the results obtained from different phosphoproteomic normalization strategies. Data were scaled, clustered and visualized as heatmap. (C) The P-site-specific spearman correlations between Method 2 and Method 3, based on the varied data among six cell lines.

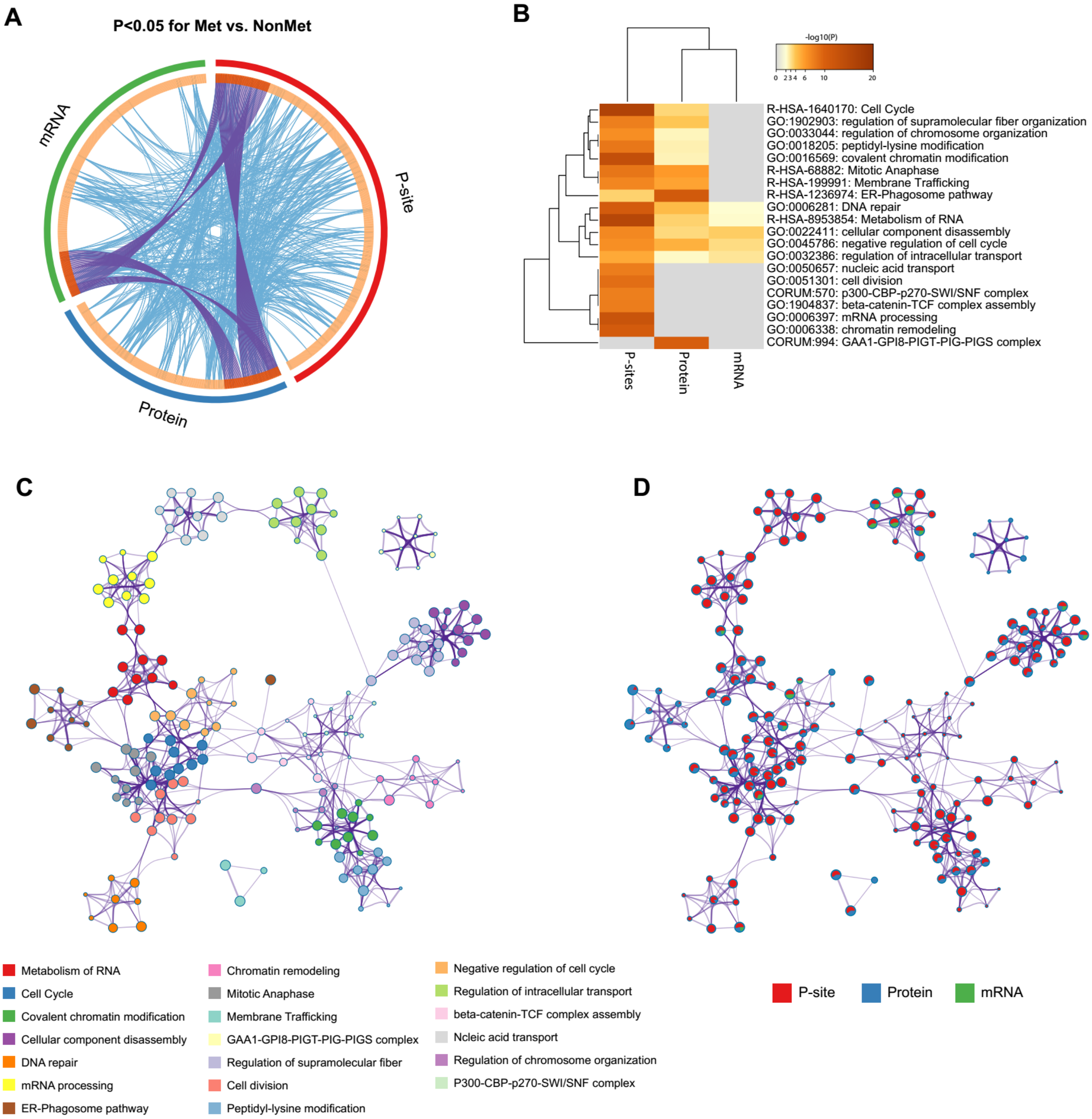


**Supplementary Figure S4 | Preliminary insights on melanoma cancer metastasis with multi-omics profiling.** (A) The circos plot showing the significant signature genes identified across the three omics layers. Each arc represents the identity of each gene list from mRNA, protein, and phosphopeptide measurements. Blue lines link the different genes where they fall into the same ontology term (the term has to statistically significantly enriched and with size no larger than 100). (B) The overlapping functional processes across the three layers. All statistically enriched terms (GO/KEGG terms, canonical pathways, and hall mark gene sets) were identified and the accumulative hypergeometric p-values calculated by Metascape were shown. (C) A subset of representative terms from the full cluster were converted into a network layout. Terms with a similarity score > 0.3 are linked. The network is visualized with Cytoscape with "force-directed" layout. (D) The same enrichment network as (C) which has its nodes colored by p-value, as shown in the legend. The dark the color, the more statistically significant the node is.

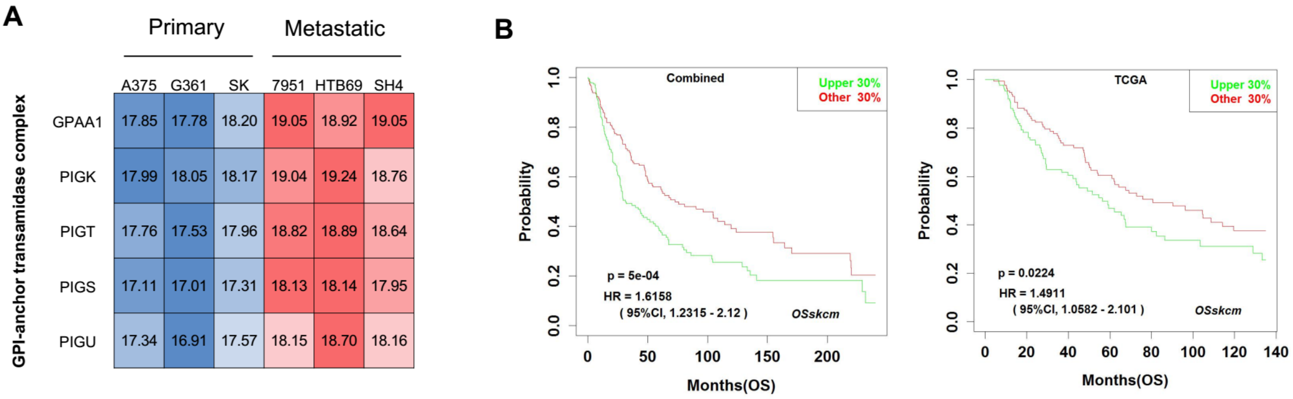


**Supplementary Figure S5 | The relationship between the overexpression of GPI-anchor transamidase complex and melanoma metastasis.** (A) The abundance of all the five subunits of this complex across cells. The log-2 values of the DIA-MS peak areas are shown, indicating a uniform ~2-fold upregulation in metastatic melanoma cells for this protein complex. (B) The predicted prognosis outcome for the mRNA levels of GPAA1 using all the clinical melanoma samples in OSskcm and TCGA datasets.
